# A Biophysical Platform for Electromechanical Stimulation of Engineered Cardiac Tissues

**DOI:** 10.64898/2026.08.31.747584

**Authors:** Ruifeng Hu, Claudia E. Varela, M. Çağatay Karakan, Francisco Sanchez, Benjamin Wolking, Christopher S. Chen, Thomas G. Bifano

## Abstract

Human engineered cardiac tissues (ECTs) provide an in vitro model for studying human cardiac physiology and drug responses, but their performance remains limited by culture systems that do not fully reproduce the heart’s electrical and mechanical environment. Electrical stimulation (ES) and mechanical stimulation (MS) have each been used to improve ECT function. Their combination, referred to as electromechanical stimulation (ES+MS), can provide further benefits. However, ES+MS depends not only on the presence of both cues but also on how they are coordinated in time. Here, we developed an incubator-compatible biophysical platform that delivers ES and MS independently or in combination, with programmable control over timing, amplitude, frequency, duration, and waveform. Calibration and dynamic characterization demonstrated tissue-relevant strain delivery, rapid and repeatable motion, and minimal attenuation and timing lag at the designated frequency of 1.5 Hz. We then compared four 6 h conditioning regimens: unstimulated control, ES alone, unsynchronized ES+MS, and synchronized ES+MS. We hypothesized that the synchronized ES+MS group, in which electrical excitation was aligned with peak externally applied strain, would produce the greatest increase in contractile force. Consistent with this hypothesis, synchronized ES+MS increased normalized twitch force by approximately 44% on average, whereas the other groups showed no comparable improvement. Twitch-timing metrics did not exhibit coordinated enhancement after 6 h, suggesting that the force increase reflects an adaptive biomechanical response rather than broad tissue maturation. These findings identify ES–MS timing as an important design parameter for ECT conditioning.

## INTRODUCTION

Engineered cardiac tissues (ECTs) are an important platform for disease modeling, drug screening, and mechanistic studies.^1–7^ Compared with two-dimensional human induced pluripotent stem cell-derived cardiomyocyte (hiPSC-CM) cultures, three-dimensional ECTs better capture tissue-level features and biomechanics of myocardium, including multicellular organization, extracellular matrix remodeling, and force generation via cell-cell coupling against defined mechanical boundary constraints.^8–13^ ECTs enable the investigation of human-specific biomechanics, electrophysiology, drug response, and disease mechanisms unable to be studied in animal models, within a controllable in vitro setting.^1,2,4,14–16^ Despite these advantages, the functional performance of ECTs still falls short of native myocardium, in part because current culture and conditioning strategies do not fully reproduce the dynamic electrical and mechanical environment of the heart.^5,7,17,18^

Electrical stimulation (ES) and mechanical stimulation (MS) have been widely explored as biophysical conditioning strategies that better mimic cardiac physiology and pathology in ECTs.^19–24^ By delivering electrical impulses at a defined rhythm, ES can control ECT excitation– contraction timing and improve pacing responsiveness, electrophysiological properties, calcium handling, sarcomere organization, and contractile performance, while supporting broader structural and functional ECT development.^5,25,26^ On the other hand, MS modulates the biomechanical loading environment of ECTs through externally applied static tension, geometric constraints, cyclic stretch, afterload-like resistance, perfusion- or shear-based loading, and more advanced pressure–volume-like paradigms.^27,28^ Both of these approaches have shown beneficial effects in ECT compaction, cardiomyocyte alignment, contractile force generation, and mechanotransduction-associated maturation.^7,24,29–33^

Previous studies have also delivered ES and MS concurrently, demonstrating that electromechanical stimulation (ES+MS) further improves ECT function when compared to either ES or MS alone.^34–36^ However, reproducing the dynamic environment of native myocardium requires more than simply applying the two inputs concurrently.^26,37,38^ In native myocardium, electrical activation and mechanical function are tightly coupled in time through excitation– contraction coupling and mechano-electric feedback, and the temporal relationship between excitation and load is a fundamental determinant of mechanical output and mechanoresponsive signaling.^39–42^ Stimulation regimes studied to date have mainly combined ES with static stretch, which does not capture the dynamic loading experienced in vivo.^35,43,44^ Fewer studies have combined ES with cyclic stretch, and in these studies, the timing between the two stimuli has generally remained fixed rather than being systematically varied to simulate diverse loading scenarios.^36,45^ Although Morgan et al. varied ES–MS timing, their rat cell–derived constructs were subjected to a square-wave stretch profile that differs substantially from the continuous deformation of the native myocardium.^37^ Thus, how the timing between ES and MS affects ECT function under dynamic loading remains unclear.

Here, we developed a biophysical platform that enables precise control over the timing between electrical and mechanical inputs, as well as stimulus amplitude, frequency, duration, and waveform. Using this system, we evaluated whether the temporal relationship between ES and MS affects ECT contractile force generation after 6 h of conditioning. Four groups were compared: unstimulated controls, ES alone, synchronized ES+MS, and unsynchronized ES+MS. In the synchronized ES+MS condition, the electrical pulse was timed to coincide with the peak of externally applied tissue strain, more closely recapitulating the physiological temporal coupling between electrical activation and mechanical loading that occurs in the native myocardium. In the unsynchronized ES+MS condition, peak strain was delayed by 200 ms relative to the electrical pulse, disrupting this relationship. We hypothesized that synchronized ES+MS would increase contractile force generation compared with unsynchronized ES+MS, ES alone, and unstimulated controls.

## RESULTS

### Biophysical Platform for ECT Stimulation

To examine how the temporal relationship between ES and MS influences ECT contractile force generation, we designed and built a custom biophysical stimulation platform. The platform can deliver ES and MS to ECTs housed in microfabricated tissue gauge devices, referred to as μTugs (Supp. Fig. 1), either independently or in combination, with the waveform of each stimulation modality programmed separately [Fig. 1(a–c)].

**Fig. 1.**
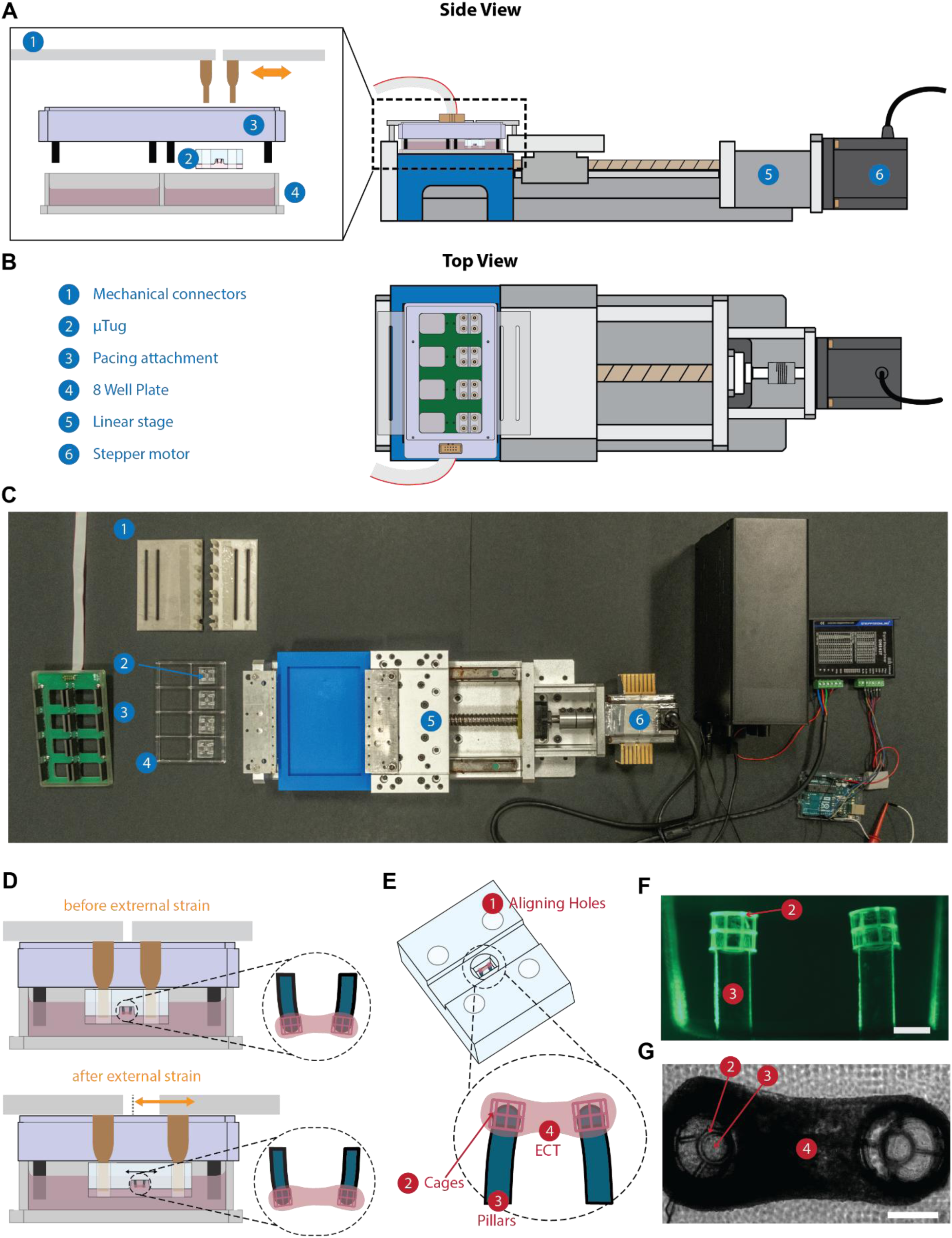
Biophysical platform for ECT conditioning. (a) Side-view schematic and enlarged view of the conditioning setup showing the ES and MS configuration. Numbered labels indicate the major system components: (1) mechanical connectors, (2) μTug, (3) pacing attachment, (4) 8-well plate, (5) linear stage, and (6) stepper motor. The same numbering scheme is used in panels (a–c). (b) Top-view schematic of the same setup. (c) Exploded-view photograph of the same setup. (d) Side-view schematic showing how displacement of the mechanical connectors stretches the μTug and applies external strain to the ECT. (e) Schematic of the μTug highlighting the central well that houses the ECT (dashed box). The same numbering scheme is used in panels (e–g). (f) Representative photograph of a μTug and fluorescence side view of the pillar pair with cages installed. Scale bars: 0.3 mm. (g) Representative top-view image of an ECT spanning the two pillars with cages in place. Scale bar: 0.3 mm. Unless otherwise noted, μTugs are shown right-side up throughout the manuscript for clarity, except in [Fig. 1(a–d)], where they are shown inverted. During conditioning and end-point measurements, μTugs were mounted inverted and submerged in culture medium.

During experiments, mechanical connectors [Fig. 1(a–c), (1)] were attached to the blocks fixed to the linear stage assembly using 4-40 screws. Each connector consists of a stainless-steel base plate fitted with four pairs of vertical prongs, each with a necked mid-section. The prongs were fabricated from biocompatible polypropylene (PP) using 3D printing and engage with the grip holes on either side of each μTug [Fig. 1(a–c), (2)], ensuring secure and reproducible positioning at a defined height. ES was delivered via a commercial pacing system (C-Pace EM, IonOptix) and an 8-well pacing attachment (C-Dish, IonOptix) with graphite electrodes [Fig. 1(a–c), (3)]. During conditioning, each μTug containing the seeded tissue was placed upside down into an 8-well plate [Fig. 1(a–c), (4)], with each well filled with 4 mL of culture medium to ensure complete submersion of the ECTs.

The pacing attachment was placed on top of the 8-well plate, with openings that allow the prongs to pass through and engage the grip holes in μTugs. A thin parafilm layer was inserted between the mechanical connectors and the pacing attachment to reduce media evaporation. The 8-well plate was rigidly fixed within a custom 3D-printed holder, shown in blue, which was seated on a linear translational ball-bearing slide [Fig. 1(a–c), (5)]. The slide was driven by a stepper motor [Fig. 1(a–c), (6)] and a ball screw to displace the mechanical connectors, thereby applying external strain to the μTugs and the ECTs housed within them [Fig. 1(d)].

The polydimethylsiloxane (PDMS) μTug device was designed to house ECTs and enable controlled mechanical stimulation based on a previously reported version.^46^ To support ES+MS, a rectangular channel was added across the center of the device, passing through the central well [Fig. 1(e)], to ensure a conductive path for the electric field across the tissue. In addition, an add-on cage structure was introduced to improve tissue fixation at the pillar tops. The cage is a cylindrical open-frame structure composed of circular rings and vertical bars, with an open bottom and a partially closed top formed by radial spokes [Fig. 1(f)]. This design provides additional attachment features for the ECT [Fig. 1(g)], reducing detachment during ES+MS.

### Characterization of Biophysical Platform for ECT Stimulation

Platform calibration was performed to establish the relationship between the applied actuator strain and measured pillar strain, providing the basis for expressing subsequent stimulation conditions in tissue-relevant terms. The actuator-imposed input (applied strain) was calculated from stage displacement measured with an optical displacement sensor (D100, Philtec) and an oscilloscope (TBS1000C, Tektronix). In contrast, pillar strain was defined as the percentage change in pillar separation and quantified by tracking the two pillars with a microscope (IX73, Olympus). Because the ECT is anchored between the two pillars, pillar strain represents the mechanical strain directly imposed on the tissue. Applied strain and pillar strain exhibited a strong linear relationship over the tested range [Fig. 2(a)]. A zero-intercept fit yielded pillar strain = 0.497 × applied strain (R² = 0.994), indicating that the pillar strain was approximately one-half of the applied strain, such that 10% of the applied strain corresponded to about 5% pillar strain.

**Fig. 2.**
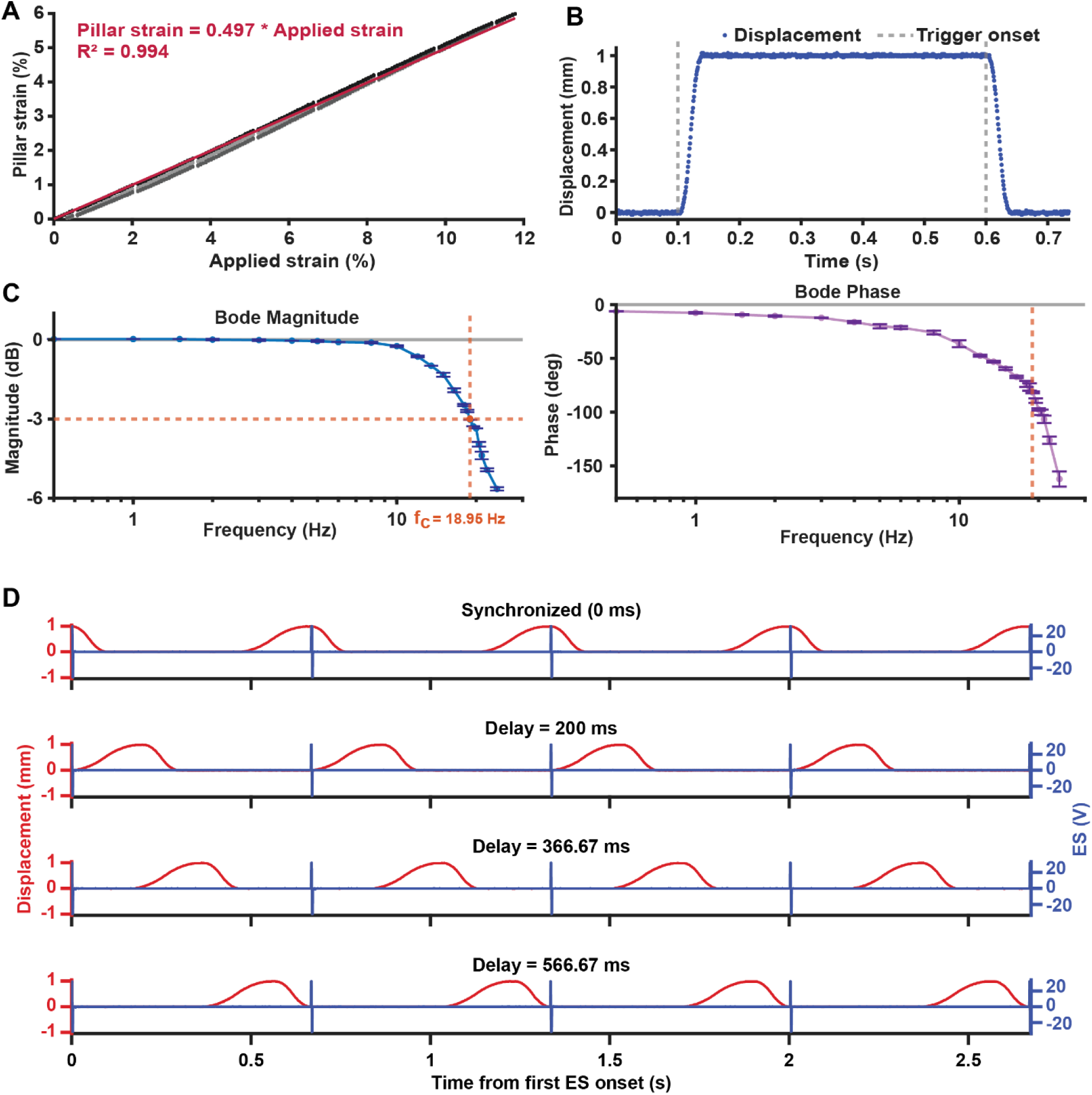
Characterization of the biophysical platform. (a) Calibration between applied strain and pillar strain. (b) Representative 1 mm step response of the system, measured under maximum achievable acceleration and deceleration conditions. (c) Bode magnitude and phase plots used to characterize frequency-dependent amplitude response and time lag of mechanical actuation. The dashed orange lines indicate the −3 dB cutoff frequency. (d) Representative measured electrical stimulation (ES) and mechanical stimulation (MS) waveforms demonstrating programmable timing between the two stimuli at 1.5 Hz. Traces show four consecutive cycles aligned to the first ES onset and ending at the completion of the fifth ES pulse. Blue traces indicate the measured electrical stimulus, and red traces indicate the measured mechanical displacement.

Next, the dynamic performance of the biophysical platform was evaluated. An alternating 1 mm step-response analysis was used to evaluate repeated reference tracking under rapid transitions [Fig. 2(b)]. Although only one representative trace is shown, measurements were repeated over 16 cycles at the highest achievable speed without slippage. The response parameters used for this characterization are illustrated in Supplementary Fig. 2, adapted from Stankevic et al., 2000.^47^ Under these conditions, the 10% delay time was 11.3 ± 0.3 ms, the 10–90% rise time was 19.7 ± 0.3 ms, and the overshoot was 0.01 ± 0.0033 mm, corresponding to 1.01 ± 0.33% for a 1 mm travel [Fig. 2(b), Supp. Fig. 2]. The measured peak velocity was 58.0 mm/s [Fig. 2(b), Supp. Fig. 2]. Together, these results indicate that the platform can execute rapid displacements with fast, repeatable, and well-controlled transient behavior.

The platform’s performance across a range of frequencies was characterized using Bode analysis. The Bode magnitude plot addresses whether the system can maintain commanded travel amplitude as frequency increases, while the phase plot quantifies the associated timing lag [Fig. 2(c)]. The magnitude response remained essentially flat at low frequencies and stayed within 1 dB through 13.5 Hz, before crossing the -3 dB cutoff at 18.95 Hz. In parallel, the phase lag increased progressively with frequency, from -6.2° at 0.5 Hz to -81.5° at 19 Hz, reaching -162.1° at 24 Hz. Notably, at 1.5 Hz, the pacing frequency used in subsequent stimulation demonstrations, the gain remained effectively unity (0.024 dB) and the phase lag was small (- 9.4°).

The main motivation for the development of this biophysical platform is to provide programmable control over the timing between ES and MS. This capability was demonstrated by recording paired ES and MS outputs under four representative timing configurations [Fig. 2(d)]. The timing conditions shown correspond to MS delays of 0, 200, 366.67, and 566.67 ms relative to an ES cycle of 666.67 ms duration (1.5 Hz stimulation frequency). Because the two outputs are generated independently from the same Arduino-based controller clock, this delay can be prescribed across the full cycle, enabling controlled comparison of synchronized and unsynchronized ES+MS conditions. In addition to the temporal relationship, the platform supports independent adjustment of stimulation amplitude, frequency, duration, and waveform for both ES and MS [Figs. S3–S5].

Taken together, these characterization results demonstrate that the biophysical platform can deliver controlled ES+MS under tissue-relevant conditions. Strain calibration established a linear relationship between actuator-level input (applied strain) and tissue-level output (pillar strain). Step-response testing confirmed rapid and well-damped transient tracking. Bode plots show minimal amplitude attenuation and timing lag at the 1.5 Hz stimulation frequency used in subsequent ECT experiments. Finally, direct measurements of paired ES and MS outputs confirmed that the two stimuli could be synchronized or deliberately offset within each stimulation cycle.

### Electromechanical Stimulation of ECTs Using the Developed Biophysical Platform

To investigate how the timing between ES and MS waveforms influences ECT function, four experimental conditions were established [Fig. 3(a, b)]. The unstimulated control group was maintained without external ES or MS, providing a functional baseline [Fig. 3(a, b), (1)]. The ES group received electrical pacing only [Fig. 3(a, b), (2)]. The unsynchronized ES+MS group received both stimuli, but the peak of externally applied cyclic strain was delayed relative to ES, reflecting a non-physiological temporal sequence of ES during active cardiac contraction [Fig. 3(a, b), (3)]. In contrast, the synchronized ES+MS group received ES at the peak of externally applied cyclic strain to simulate electrical activation under increased cardiomyocyte preload at end diastole [Fig. 3(a, b), (4)]. Together, these groups allowed timing-specific effects of ES+MS to be distinguished from the general effects of ES or culture alone.

**Figure 3.**
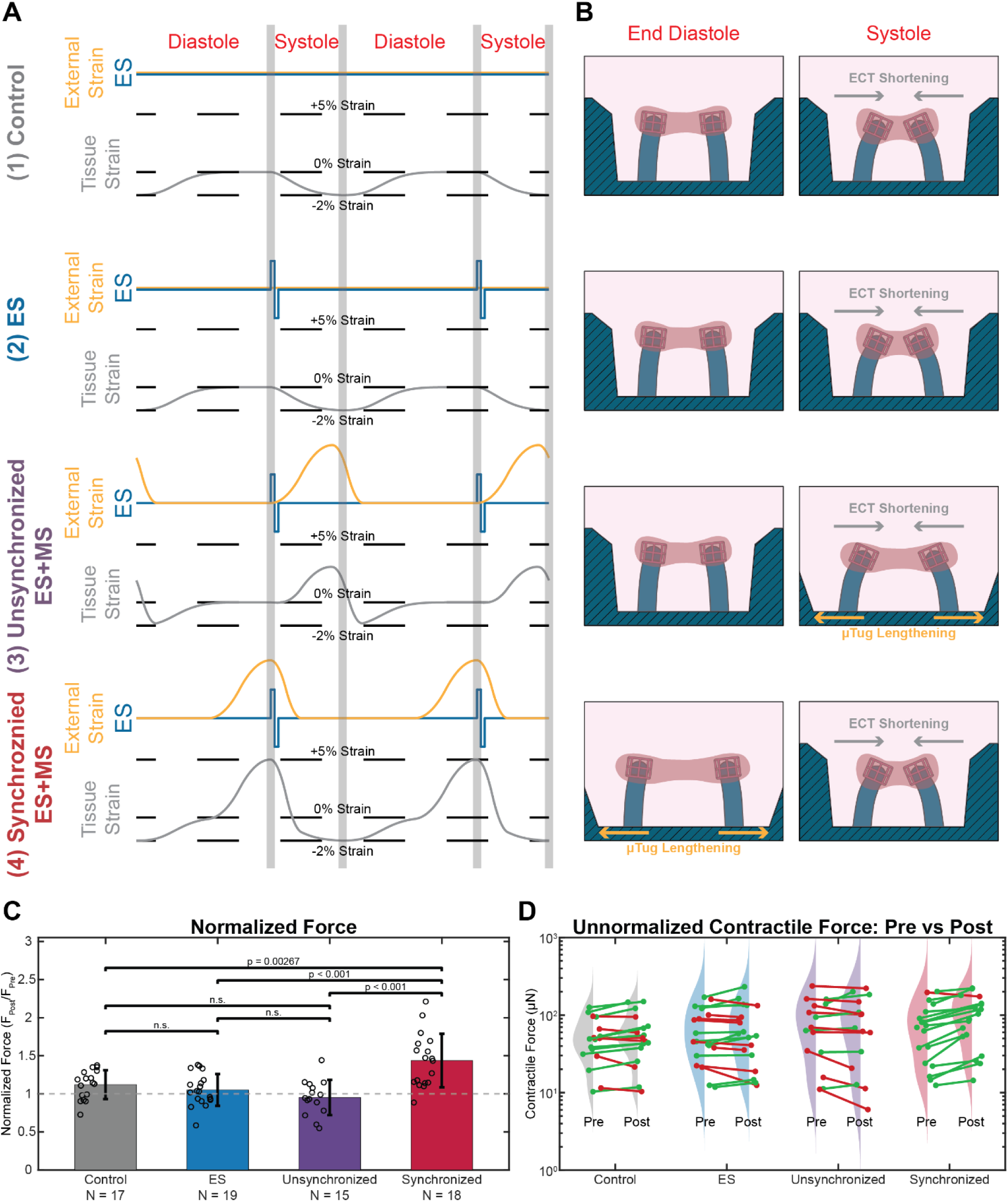
Experimental conditions and contractile force response after 6 h conditioning. (a, b) Schematic overview of the four experimental groups and the corresponding timing relationships between externally applied strain and ES. Rows (1)-(4) correspond to Control, ES, Unsynchronized ES+MS, and Synchronized ES+MS, respectively. Orange traces/arrows indicate externally applied cyclic strain, blue traces indicate ES, and gray traces/arrows indicate tissue strain/deformation. Tissue strain refers to the total strain experienced by the ECT. In the Control and ES groups, where no external MS was applied, tissue strain was entirely tissue-generated, either spontaneous or ES-evoked. In the ES+MS groups, tissue strain represents the combined contribution of tissue-generated strain and externally applied cyclic strain. (c) Post/Pre normalized twitch force after 6 h conditioning. Bars represent group means, points represent individual samples, and error bars indicate standard deviation. The dashed line at *F_Post_* / *F_Pre_* = 1 indicates no change after conditioning; horizontal brackets indicate statistical comparisons. (d) Absolute twitch force measured before and after conditioning for each sample. Paired points are connected by lines, with green indicating increased force after conditioning and red indicating no change or decrease.

Contractile force generation is a primary functional readout for ECT, as it directly reflects the tissue’s ability to generate twitch contractions. For each sample, contractility was assessed immediately before conditioning (Pre) and after 6 h of conditioning (Post). During both measurements, all samples were electrically paced at 1.5 Hz to standardize the measurement conditions. Sample sizes were N=17 (Control), N=19 (ES), N=15 (Unsynchronized ES+MS), and N=18 (Synchronized ES+MS). To decouple variations in inter-sample baseline contractility from variations in tissue contractility attributable to conditioning, post-conditioning force was normalized by pre-conditioning force for all samples [Fig. 3(c)]. Absolute (unnormalized) contractile forces are shown alongside the normalized data to display the underlying pre- and post-conditioning distributions and the paired, within-sample changes [Fig. 3(d)].

After 6 hours of conditioning, the synchronized ES+MS group showed a 44% increase in normalized contractile force on average [Fig. 3(c)]. In contrast, the ES and unsynchronized ES+MS groups were not significantly different from the Control group, and they were also not significantly different from each other. Together, these results indicate that the timing between ES and MS has an impact on ECT contractility within 6 h; neither ES alone nor unsynchronized ES+MS produced a statistically significant increase in force over the same 6 h conditioning window. Notably, whereas prior studies typically require days to weeks of electrical pacing or cyclic stretch to detect functional gains,^5,43,48^ we observed a measurable increase in force within hours when the two stimuli were applied in a physiologically synchronized manner.

To assess whether conditioning altered twitch kinetics, we quantified timing metrics that capture the relative rates of contraction and relaxation and provide a compact description of twitch shape (e.g., sharper versus more prolonged responses) before and after conditioning (TTP, CT50, CT90, RT50, RT90, and TD50) [Fig. S6]. For contraction-phase metrics, TTP increased in ES and was unchanged elsewhere [Fig. 4(a)]. CT50 decreased in Control and Synchronized and was unchanged in the remaining groups [Fig. 4(b)]. CT90 decreased in Control but increased in ES, with no detectable change in the other groups [Fig. 4(c)]. For relaxation-phase metrics, RT50 increased in Control, ES, and Unsynchronized [Fig. 4(d)], while RT90 increased in ES and Unsynchronized [Fig. 4(e)]. Finally, TD50 increased in ES and Unsynchronized and was unchanged in Control and Synchronized [Fig. 4(f)].

**Figure 4.**
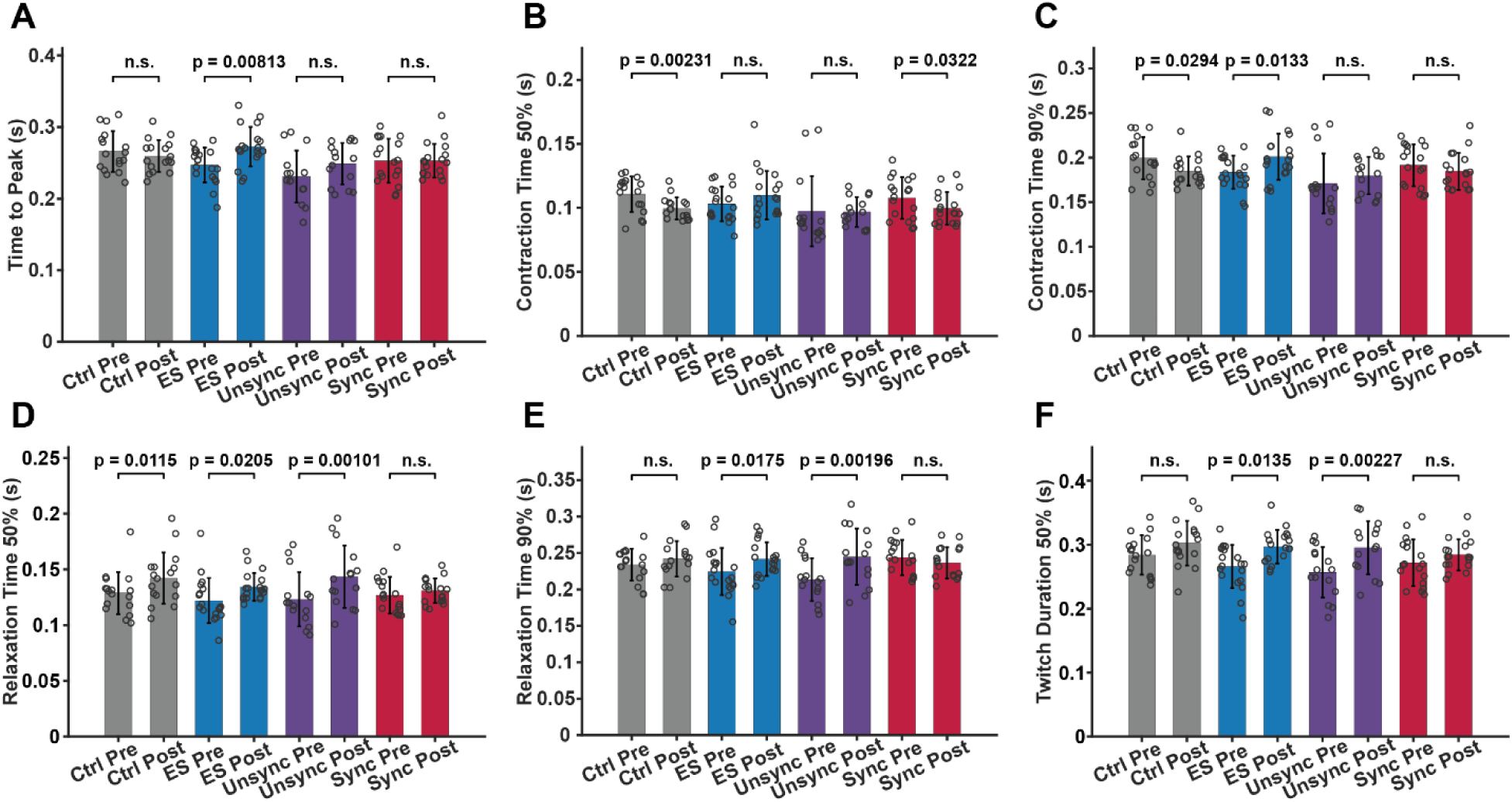
Changes in twitch-timing metrics after 6 h conditioning. Twitch-timing metrics were quantified before (Pre) and after (Post) conditioning for Control, ES, Unsynchronized ES+MS, and Synchronized ES+MS groups. Metrics include (A) time to peak (TTP), (B) contraction time 50% (CT50), (C) contraction time 90% (CT90), (D) relaxation time 50% (RT50), (E) relaxation time 90% (RT90), and (F) twitch duration 50% (TD50), as defined in Fig. S6. Bars represent group means, points represent individual samples, and error bars indicate standard deviation. Horizontal brackets indicate within-group Pre–Post comparisons, with exact p-values shown where significant and n.s. indicating no significant difference.

## DISCUSSION

In this work, we developed and validated a biophysical platform that can generate electromechanical stimulation for ECTs, featuring the ability to program ES and MS individually and precisely. The system was designed to deliver independently specified electrical and mechanical waveforms in an incubator-compatible format. Platform characterization validated that ES and MS timing could be imposed with minimal amplitude attenuation and timing lag at the 1.5 Hz stimulation frequency used for ECT conditioning. Using this platform, we demonstrate that the temporal relationship between ES and MS is a key determinant of the effectiveness of ECT electromechanical conditioning.

The synchronized ES+MS stimulation protocol produced a substantial (44%) enhancement in ECT contractile function as compared to any other protocol in this study. Importantly, unsynchronized ES+MS, despite employing identical electrical and mechanical waveforms, failed to elicit comparable improvements and was functionally indistinguishable from either ES alone or control conditions. These results align with prior reports that ES+MS improves ECT function more than either cue alone,^34–37,43^ but highlight that the presence of dual stimulation itself is not sufficient to elicit a functional change, and the timing between the two stimuli is critical. This may help interpret earlier reports in which simultaneous dual stimulation was beneficial, but the specific contribution of timing was unclear.^34–37,43^

In vivo, the contractile force generated by the myocardium increases with venous return or preload (i.e., sarcomere length just before contraction), as described by the Frank-Starling mechanism. Our synchronized ES+MS group was designed to harness this mechanism by aligning peak stretch with electrical excitation, thereby increasing ECT preload with respect to the other experimental groups. The enhanced ECT contractile function observed after only 6 h of synchronized conditioning indicates that ECTs are sensitive to this physiologically relevant loading and suggests that loading state, controlled by ES+MS timing, may be an important parameter to investigate in longer-term electromechanical stimulation regimes designed to improve ECT functional maturation.

In contrast to the force response, ES+MS did not yield notable changes in twitch-timing metrics after 6 h of conditioning. Improved twitch kinetics are closely linked to more efficient calcium cycling and electrophysiological properties,^49,50^ which are commonly evaluated as part of longer-term (weeks) ECT maturation studies.^5,25,43,48^ Thus, the increased contractile force observed in the synchronized ES+MS group should be interpreted as a relatively rapid adaptive biomechanical response rather than evidence of broad maturation.

Beyond the biological finding, this study showcases the value of a modular stimulation platform for testing electromechanical timing as an independent experimental variable. The platform’s compact footprint allows stimulation to occur entirely inside a standard commercial incubator, avoiding custom environmental chambers that increase complexity and reduce repeatability.^51,52^ Aside from the mechanical connectors, the system is built largely from readily available commercial components. For actuation, we prioritized delivering the same uniaxial strain waveform economically and with high positional precision using a stepper-motor-driven linear stage. This choice contrasts with more specialized approaches, such as magnetic actuation, pneumatic deformation of elastomeric structures, or external actuator and force sensor transfer rigs, which can be effective but often add complexity, slow response, or cumbersome sample handling.^30,53–57^ Additionally, the modular platform architecture makes it straightforward to substitute components to fit other laboratories’ setups, promoting adoptability.

Because this study focuses on the timing between ES and MS, stimulation parameters were carefully selected to minimize confounding variables that could obscure timing-dependent effects. For ES, threshold-based biphasic pacing was used to support reliable ECT capture while limiting unnecessary electrical exposure and electrode–electrolyte reactions. For MS, dynamic cyclic stretch was selected over static stretch, as prior studies suggest that cyclic loading provides greater functional benefit within a physiological strain range.^34,58,59^ Given that higher strains, such as 10–20% at approximately 1 Hz, may impair function or induce tissue damage,^60^ we used a moderate 5% strain amplitude to avoid masking timing effects, driving pathological remodeling, or depressing contractile performance.^61–63^ Since cardiac mechano-electric signaling and tissue mechanics depend on strain rate as well as amplitude, we used a smooth asymmetric piecewise-parabolic mechanical stimulation waveform rather than abrupt triangular or trapezoidal strain waveforms that can introduce rate-dependent effects complicating comparisons across ES+MS timing conditions.^64,65^ Lastly, we did not attempt to reproduce isovolumetric contraction or relaxation as strict boundary conditions, although some conditioning studies have closely mimicked the Wiggers diagram with isovolumic “dead zones” in the loading waveform,^53,54^ because ECTs lack valves or a pressure–volume chamber.

Several limitations remain. First, our study isolates timing effects at a fixed frequency (1.5 Hz) rather than addressing frequency as an additional variable. Second, our conditioning window was long enough to detect short-term effects, but it is unlikely to capture longer-term ECT maturation processes such as sarcomere reorganization, metabolic remodeling, or T-tubule development, which typically require days to weeks of conditioning.^5,43,48^ Third, our readouts were primarily mechanical and kinematic, including force and twitch-timing metrics extracted from pillar deflection. We did not directly measure electrophysiological parameters such as calcium handling or membrane voltage, limiting mechanistic attribution related to excitation–contraction coupling or mechano-electric feedback. Finally, generalizability is constrained by the use of a single hiPSC line, a fixed CM ratio, and the specific ECT geometry used here.

Building on these limitations, future work using the platform presented here will extend conditioning to multi-day and week-long regimens to determine if the improved ECT contractility after synchronized ES+MS persists or diminishes over time. Additional studies can systematically vary stimulation frequency, electrical field strength, strain amplitude, waveform shape, and ES–MS timing to investigate the biological effects of various electromechanical ECT conditioning regimes. Frequency ramp protocols (e.g., 1 to 6 Hz in daily 1 Hz increments) have been reported to improve engineered cardiac tissue function and promote maturation over longer periods.^20,25,66^ In fact, to our knowledge, ramping of both electrical and mechanical stimulation has not been explored. To strengthen mechanistic interpretation, future studies should incorporate additional readouts, including calcium transients, structural assays of sarcomeres and T-tubules, and molecular or metabolic profiling. Finally, integrating an add-on imaging module into the incubator-ready platform would enable real-time tracking during conditioning rather than relying solely on endpoint measurements, and expanding to multiple hiPSC lines would help assess translational relevance.

Overall, this study shows that the temporal relationship between ES and MS is an important design parameter for ECT conditioning, and highlights the utility of our programmable platform for systematically investigating these effects. More broadly, these findings suggest that improving the physiological relevance of engineered cardiac tissue platforms requires not only applying electrical and mechanical cues, but also controlling how those cues are coordinated in time.

## METHODS

### Elastomeric μTug Fabrication

The PDMS μTug was fabricated following the mold-casting protocol outlined in our previous work.^46^ Molds were initially designed in CATIA and produced via stereolithography (Protolabs) using MicroFine Green resin (layer thickness = 25 μm; minimum supported feature size = 70 μm). To minimize adhesion during demolding, the received printed components underwent vapor-silanization. During casting, the 3D-printed molds were filled with a Sylgard 184 Silicone Elastomer mixture (monomer to crosslinker ratio of 10:1), then cured overnight on a 60 °C hot plate [Fig. 5(a)].

**Fig. 5.**
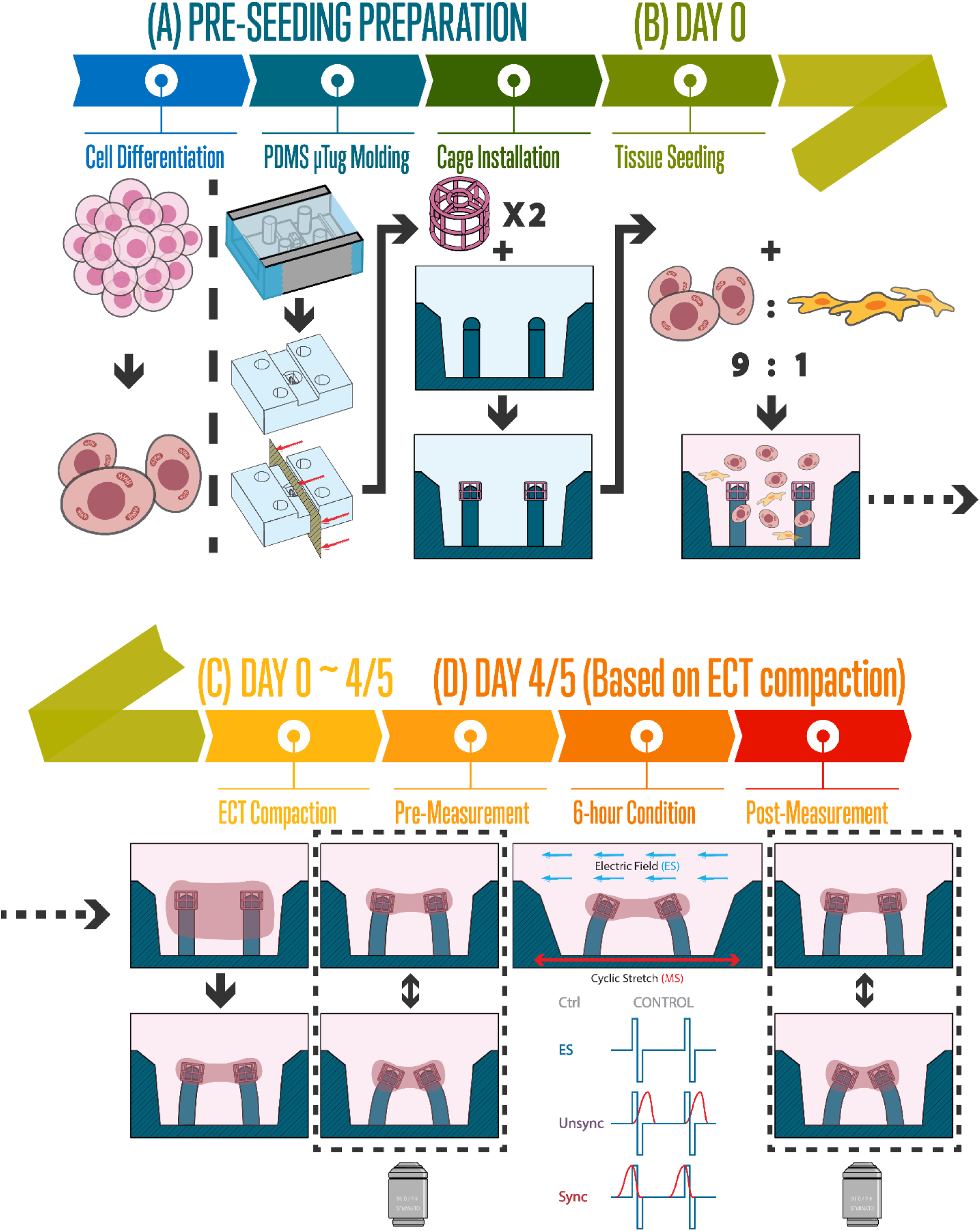
Experimental workflow for ECT preparation, conditioning, and endpoint measurements. (a) Pre-seeding preparation: hiPSCs were differentiated into cardiomyocytes in parallel with fabrication of elastomeric PDMS μTug devices by mold casting. After demolding, cage add-ons (two per μTug; one per compliant pillar) were installed to provide defined tissue attachment sites at the pillar tops. (b) Day 0 (seeding): cardiomyocytes were mixed with cardiac fibroblasts at a 9:1 ratio and seeded into each μTug (60,000 total cells per μTug). (c) Days 0–4/5: most seeded tissue reached a stable compacted morphology by day 4, with some requiring an additional day. (d) Day 4/5 (based on completion of compaction): a pre-conditioning endpoint measurement (“Pre”) was performed, followed by a 6-h conditioning period under one of four regimens: Control, ES, Unsync (unsynchronized ES+MS stimulation), or Sync (synchronized ES+MS stimulation). A second endpoint measurement (“Post”) was then performed for all groups.

The contractile force of the ECT was quantified by tracking the bending of the two flexible pillars. The effective pillar stiffness (k) was measured as k = 4.82 ± 0.013 N/m using micro-indentation.^46^ The pixel-to-length conversion for the 2× and 4× objectives was calibrated using a stage micrometer (Thorlabs), yielding values of 5.676 µm/pixel and 2.838 µm/pixel, respectively. These calibration factors were used to convert the tracked pillar displacements (δ) into force (F) using Hooke’s law (F = kδ). In this analysis, the deflections of the two pillars were equal in magnitude but opposite in direction.

### Fabrication and Assembly of Tissue Attachment Sites

We initially observed low yield in experiments due to the detachment of the ECTs from the PDMS pillars, and high variance in force measurements, which could be due to the variable positioning of the ECTs along the pillar axis. We attempted to address these issues by fabricating and assembling customized attachment sites on PDMS pillars [Fig. 5(a)]. These attachment sites [Fig. 1(e–g)], which we refer to as “cages”, are porous cylinders that support localized tissue attachment and stability, based on our previous works.^67,68^ We fabricated a 10×10 array of these cages via a commercial two-photon direct laser writing system (GT2, Nanoscribe Professional), using a 25× objective (NA=0.8) in a dip-in mode to scan the laser (slicing distance=1.8 µm, hatching distance=0.8 µm, 100 mW laser power, 50 mm/s scanning speed) over the IP-S resin on a silicon substrate. After printing, non-polymerized resin was rinsed by immersing the substrate vertically in a propylene glycol methyl ether acetate (PGMEA) bath for 20 min, followed by isopropanol for 1 min, and NOVEC 7000 (3M) for 1 min to remove residual solvent and dry the cages. The resulting cages were approximately 500 µm in diameter and 420 µm in height, with scaffolding elements 40 µm in diameter. Next, we inserted the cages on pillars using a tweezer under a stereo microscope (MZ7.5, Leica). A representative image after cage assembly on pillars is provided [Fig. 1(f)], in which cages exhibit fluorescence in green when excited with blue light (TE200, Nikon). To improve the adhesion of cages to PDMS pillars, we plasma-treated the assemblies for two minutes (PDC-001, Harrick Plasma), immediately followed by 1-2 h on an 80 °C hot plate, typically within 24 hours prior to tissue seeding. After adopting this methodology and using a surfactant (2% Pluronics F-127) to reduce unwanted cell/protein adhesion to PDMS, we consistently observed tissue formation and attachment around the cages assembled on top of the pillars. This methodology improved our yield by 4× compared to the condition with bare PDMS pillars (based on 9 days after seeding) and provided consistent tissue positioning on the pillar tops, which reduces the variability in stiffness (i.e., afterload) ECTs experience and makes the force measurements more accurate.

### Cell Sources and Differentiation

Cell maintenance and cardiomyocyte differentiation followed our previously published methods.^69,70^ Briefly, we used a PGP1 human iPSC line (Personal Genome Project) carrying an endogenous titin–GFP reporter generated by CRISPR/Cas9 editing.^71^ Differentiation was initiated once cultures reached ∼90% confluence using a monolayer WNT modulation protocol in RPMI 1640 (Gibco) supplemented with B27 (Thermo Fisher Scientific) minus insulin and GlutaMAX (Fisher).^69^ WNT signaling was activated on day 0 with CHIR99021 (Tocris, 12 μM, 24 h) and subsequently inhibited on day 3 with IWP4 (Tocris, 5 μM, 48 h). From day 7 onward, cells were maintained in RPMI containing B27 with insulin. After the onset of spontaneous contractions, cardiomyocytes were enriched by metabolic selection using sodium lactate (Sigma, 4 mM, 4 d), replated on fibronectin-coated plates, and evaluated by flow cytometry, yielding 81.1 ± 1.9% viability and 94.6 ± 1.4% cTnT⁺ purity [Fig. 5(a)].^70^

### Engineered Cardiac Tissue Seeding

Tissue Seeding process is essentially embedding 54,000 iPSC-CMs and 6,000 cardiac fibroblasts (CFs) in a scaffold made of Matrigel and fibrin (formed by fibrinogen and thrombin). Devices were sterilized with 70% ethanol (Thermo Fisher Scientific) and UV exposure. Base of the seeding wells were coated with 2.5 µL of 5% Pluronic F-127 (Thermo Fisher Scientific) for 30 min at room temperature, then rinsed off and fully air-dried. Cells were seeded at 60,000 per well with a 9:1 ratio between iPSC-CMs and CFs. All components were kept on ice to limit premature gelation. Thrombin was prepared in H1 medium, which is made of high-glucose DMEM (Corning) + 10% FBS (Gibco) + 1% Penstrep (Gibco) + 1% NEAA (Gibco) + 1% GlutaMAX (Gibco), at 0.6 U thrombin (Fisher) per mg fibrinogen (Sigma). CFs and iPSC-CMs were dissociated, counted, combined, centrifuged, and resuspended (RPMI + 5 µM Y-27632) before mixing with Matrigel and fibrinogen (no bubbles) and seeding. Constructs were cultured in H1 + aprotinin (0.033 mg/mL; half on day 7; Sigma) with Y-27632 (Sigma) for the first 2 days, changing media every other day and adding PBS around devices to reduce evaporation [Fig. 5(b)]. Following seeding, samples underwent a 96-hour compaction period prior to baseline recordings. If the tissue sample’s degree of compaction was suboptimal, the sample was allowed an additional 24 hours for adequate compaction (this situation was rarely encountered) [Fig. 5(c)].

### Electrical Stimulation

Electrical stimulation (ES) was delivered using charge-balanced biphasic square-wave pulses at an electric field strength of 10 V/cm, a pulse duration of 4 ms, and a repetition rate of 1.5 Hz [Fig. 5(d)]. The field strength was selected to be approximately 1.5 times the measured excitation threshold, providing reliable tissue capture while avoiding unnecessarily high stimulation amplitudes.^72–76^ The electrode plates were spaced 3.17 cm apart; therefore, a 32 V potential was applied across each electrode pair to generate the desired field strength (32 V / 3.17 cm = 10 V/cm). Each stimulus pulse had a total duration of 4 ms, split evenly between a 2 ms positive phase and a 2 ms negative phase, because 2–4 ms pulses can minimize electrolysis while supporting effective pacing.^77,78^ Stimulation was delivered at 1.5 Hz, consistent with the target contraction frequency used throughout the study.

ES was applied using a C-Pace EM pacemaker (IonOptix), which was externally triggered by an Arduino Uno microcontroller (Arduino) and delivered ES through a C-Dish pacing attachment (IonOptix) equipped with parallel-plate graphite electrodes. The parallel-plate configuration was selected to provide a spatially uniform electric field across the culture well, reducing stimulation heterogeneity that can occur with rod-type electrodes. Graphite electrodes were used because of their stability and biocompatibility during prolonged ES. In preliminary testing, metallic electrodes, including gold and silver, exhibited visible electrolysis and electrode degradation and were therefore discontinued. To assess whether chronic ES introduced appreciable heating, the temperature of culture medium was monitored during a 12 h continuous pacing test, twice the experimental conditioning duration, under identical electrical settings; temperature increased only from 37.2 to 37.3 °C over the test period.

### Mechanical Stimulation

Mechanical stimulation (MS) was applied as cyclic uniaxial stretch with a 5% pillar strain, a loading duration of 200 ms, an unloading duration of 100 ms, and a repetition rate of 1.5 Hz [Fig. 5(d)]. Each cycle consisted of a smooth asymmetric strain pulse generated by a triangular velocity profile, in which the tissue was stretched from 0% to 5% pillar strain during the loading phase and then released from 5% to 0% pillar strain during the unloading phase, followed by a no-stretch interval before the next cycle. This piecewise constant-acceleration trajectory produced a continuous, piecewise-parabolic strain waveform and avoided abrupt step changes in strain. Because deformation of the compliant PDMS μTug structure does not translate directly to tissue deformation, calibration between μTug strain and pillar strain is described in the Results section. A μTug strain of approximately 10% was required to generate a tissue-level, or pillar, strain of approximately 5%. Given the 10 mm resting separation between the μTug grip holes, this condition was achieved by increasing the grip-hole spacing to 11 mm during loading, corresponding to a 1 mm displacement and 10% μTug strain.

Mechanical loading was generated by a P Series IP65 waterproof NEMA 23 stepper motor (OMC Corporation Limited) mounted to a linear rail guide slide stage (model HRC150-200B2, SFU1605 ball screw, HG15 linear guides; Heechoo), which converted motor rotation into controlled linear translation. The SFU1605 ball screw has a 5 mm lead, corresponding to 5 mm of linear travel per motor revolution. The motor was driven by a DM542T stepper driver (OMC Corporation Limited) configured at 400 pulses per revolution, yielding 1 mm of travel per 80 steps (5 mm/rev ÷ 400 pulses/rev). The same Arduino Uno microcontroller (Arduino) used to control ES generated the prescribed displacement waveform for MS, enabling control of the temporal relationship between these two stimuli.

### Environmental Control

The compaction and conditioning were performed in a HERACELL VIOS 160i CO₂ incubator (Thermo Fisher Scientific) maintained at 37 °C, 5% CO₂, and 95% relative humidity [Fig. 5(d)]. Culture medium was exchanged every other day.

### Measurement Environment and Imaging Acquisition

For both pre-conditioning and post-conditioning measurements, samples were transferred to a 35 mm Petri dish and imaged on a Research Inverted Microscope IX73 (Olympus). During recordings, temperature was maintained at 37 °C with tape heaters mounted around the dish. Humidity control was not required because the measured μTug device was fully submerged in 4 mL of culture medium within the Petri dish. CO₂ regulation was also unnecessary during measurements because each recording session was brief (less than 2 min), minimizing the risk of measurable pH drift. Each session included approximately 1 min for positioning the μTug under the microscope, 40 s for the ECT to reach steady state, and 15 s for data acquisition. Brightfield imaging was performed at 500 Hz using a 4×/0.16 objective (Olympus) with an exposure time of 1.5 ms. The field of view was set to 0.363 mm × 11.624 mm (64 × 2048 pixels). It was intentionally selected as a narrow strip to increase readout speed and achieve the target frame rate. Image sequences were acquired with a pco.edge 4.2 CLHS sCMOS camera (Excelitas Technologies Corp.), and 7000 frames were recorded per sample. During imaging, electrical pacing was delivered throughout the measurement session using graphite electrodes (AR-14, 0.0984 × 0.3940 × 0.5510 inches; Ohio Carbon Blank) to generate a uniform electric field of 10 V/cm.

### Image Processing and Parameter Extraction

The top positions of both pillars, corresponding to both ends of the tissue sample, were tracked using normalized 2D cross-correlation with sub-pixel localization obtained by a 2D quadratic fit around the correlation peak. Although displacement of the pillars is expected to occur predominantly along the x-axis and motion in the y-axis is minimal, the tracking template height was set to 0.8× the image height to enable monitoring of Cartesian displacement and to identify any non-negligible y-direction drift. Recordings showing appreciable y-direction drift were not used for analysis; the corresponding sample was remeasured. To improve robustness, measured displacements were referenced to the well edges to correct for stage drift (a nearly negligible occurrence). Displacements were then converted from pixels to tissue length (µm) using the calibrated pixel-to-length conversion, yielding a tissue length versus time trace from which contractile force and timing metrics were extracted.

### Statistical Analysis

Samples classified as uncompacted, defined as generating peak twitch force <10 µN during the pre-conditioning measurement, were excluded prior to statistical testing. For contractile force, group differences were evaluated using a one-way ANOVA followed by a Tukey–Kramer post hoc test to perform all pairwise comparisons among the four conditioning groups. For timing metrics, pre- and post-conditioning values were compared within each group using paired t-tests to assess whether conditioning altered twitch kinetics relative to each sample’s own baseline.

## SUPPLEMENTARY MATERIAL

See the supplementary material for additional details on the CAD geometry and dimensions of the PDMS μTug (Fig. S1), definitions of the step-response performance metrics used for platform characterization (Fig. S2), programmable control of mechanical stimulation amplitude (Fig. S3), duration (Fig. S4), and waveform (Fig. S5), and schematic definitions of the twitch-timing metrics used to assess contraction and relaxation kinetics (Fig. S6).

## ACKNOWLEDGEMENTS

This work was supported by National Science Foundation Engineering Research Center on Cellular Metamaterials grant EEC-1647837. Claudia E. Varela acknowledges financial support from the Ford Foundation Postdoctoral Fellowship and National Institutes of Health grant # T32HL007572.

## AUTHOR DECLARATIONS

### Ethics Approval

Ethics approval is not required.

### Author Contributions

**Ruifeng Hu:** Conceptualization (equal); Methodology (equal); Software (lead); Investigation (equal); Formal Analysis (lead); Data Curation (lead); Visualization (lead); Writing – Original Draft (lead); Writing – Review & Editing (equal). **Claudia E. Varela:** Methodology (equal); Investigation (equal); Formal Analysis (supporting); Visualization (supporting); Writing – Original Draft (supporting); Writing – Review & Editing (equal). **M. Çağatay Karakan:** Investigation (equal); Writing – Original Draft (supporting). **Francisco Sanchez:** Investigation (supporting). **Benjamin Wolking:** Investigation (supporting). **Christopher S. Chen:** Conceptualization (equal); Funding Acquisition (lead); Resources (equal); Project Administration (supporting); Supervision (supporting); Writing – Review & Editing (equal). **Thomas G. Bifano:** Software (supporting); Funding Acquisition (supporting); Resources (equal); Project Administration (lead); Supervision (lead); Writing – Review & Editing (equal)

## DATA AVAILABILITY

The data that support the findings of this study are available from the corresponding author upon reasonable request.

