## Supplementary Information for "A Biophysical Platform for Electromechanical Stimulation of Engineered Cardiac Tissues"

### SUPPLEMENTARY MATERIAL

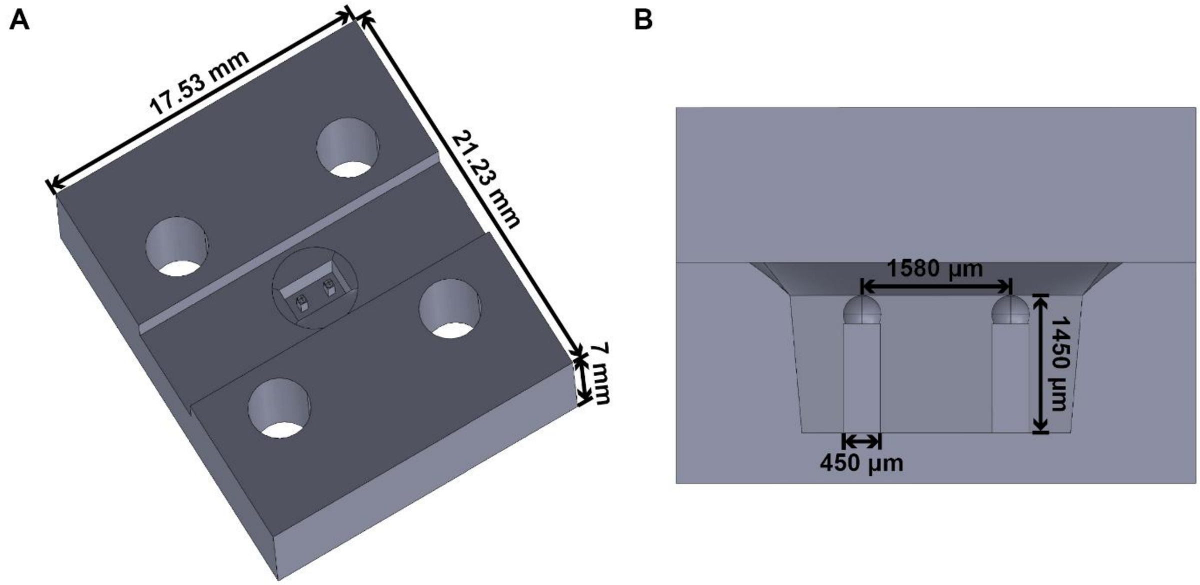


Figure S1. CAD rendering of the PDMS µTug design. (A) Isometric view showing the overall µTug geometry, central tissue chamber, and four grip holes used to interface with the stretching assembly. The µTug body measures 17.53 mm × 21.23 mm × 7 mm. (B) Cross-sectional view of the central well showing two compliant pillars that anchor the ECT, with each pillar measuring 1450 µm in height and 450 µm in width and separated by 1580 µm.

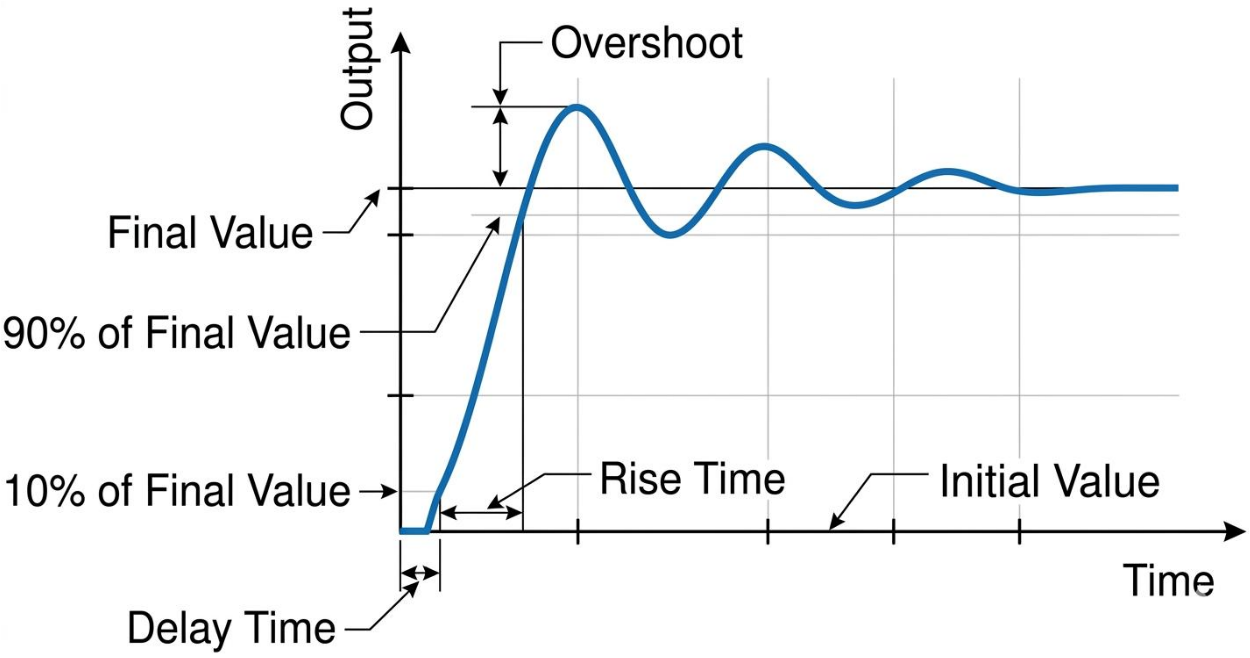


Figure S2. Definition of performance benchmarks used in this study. A schematic showing 10% delay time, 10–90% rise time, and overshoot. Adapted from Stankevic et al., 2000.[47]

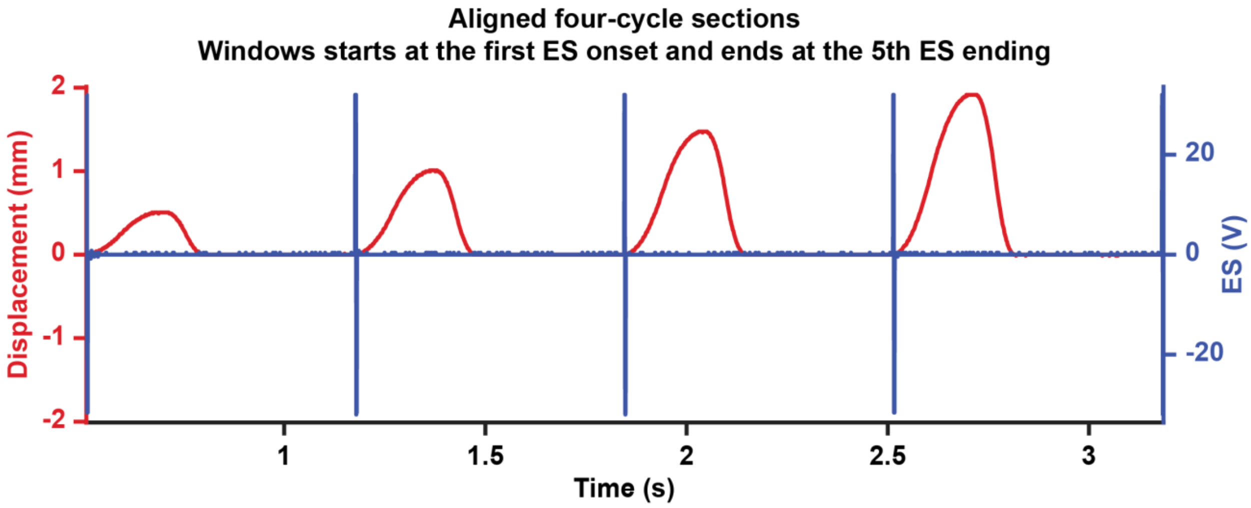


Fig. S3. Representative measured ES+MS output recordings demonstrating programmable control of mechanical stimulation amplitude. ES was kept constant, while actuator displacement was increased across successive cycles to produce travel distances of 0.5, 1.0, 1.5, and 2.0 mm. Blue traces indicate measured electrical stimuli, and red traces indicate measured mechanical displacement.

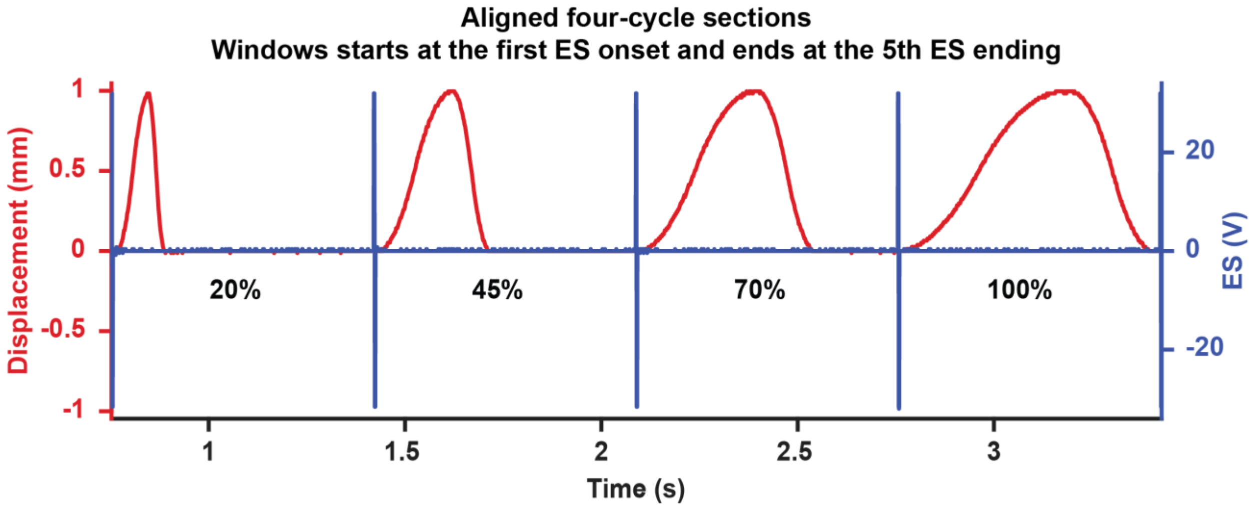


Fig. S4. Representative measured ES+MS output recordings demonstrating programmable control of mechanical stimulation duration. ES was kept constant, while the duration of actuator displacement was increased across successive cycles to 20%, 45%, 70%, and 100% of the stimulation period. Blue traces indicate measured electrical stimuli, and red traces indicate measured mechanical displacement.

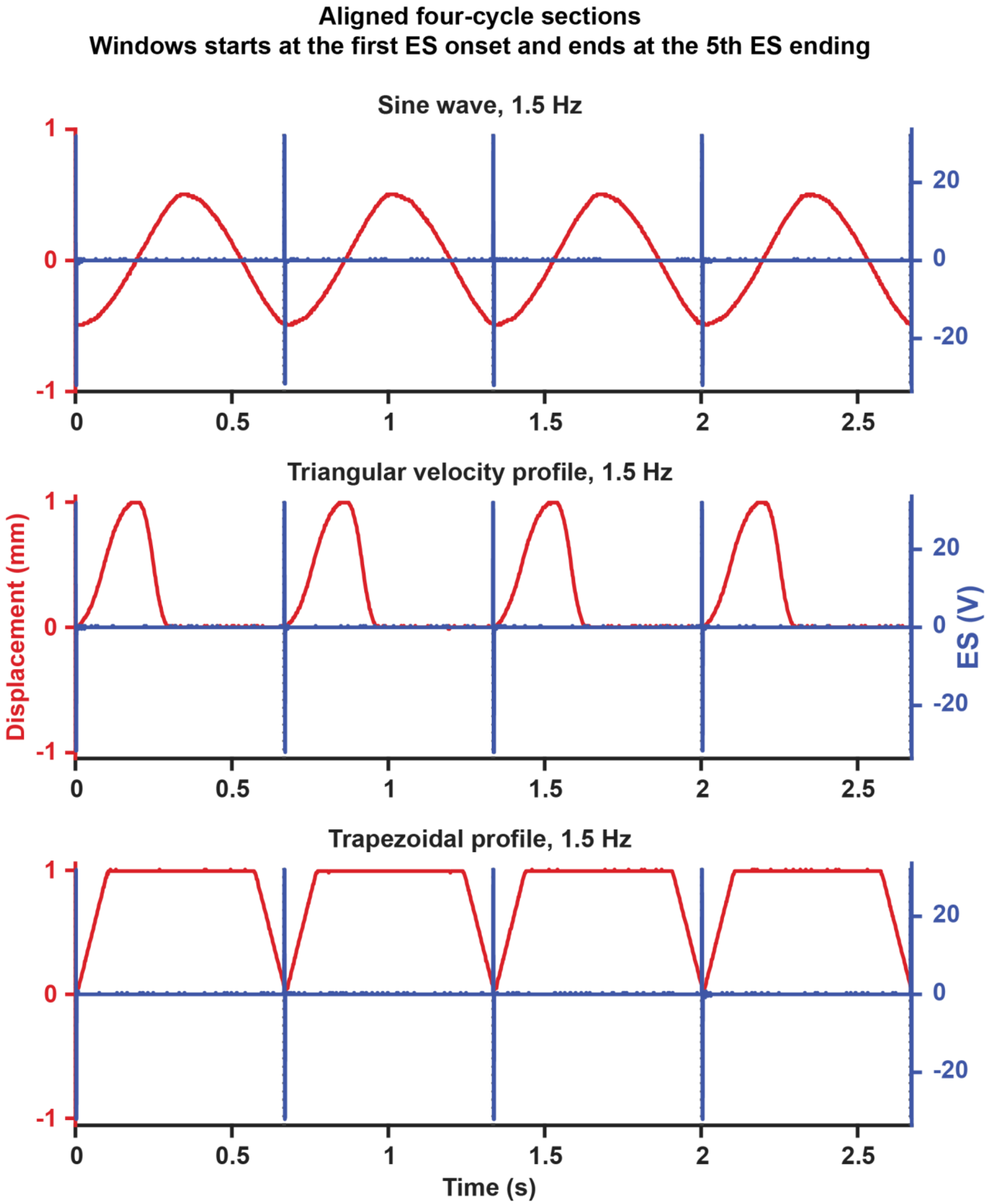


Fig. S5. Representative measured ES+MS output recordings demonstrating programmable control of mechanical stimulation waveform. ES was kept constant, while the actuator displacement profile was varied across sine-wave, triangular-velocity, and trapezoidal waveforms at 1.5 Hz. Blue traces indicate measured electrical stimuli, and red traces indicate measured mechanical displacement.

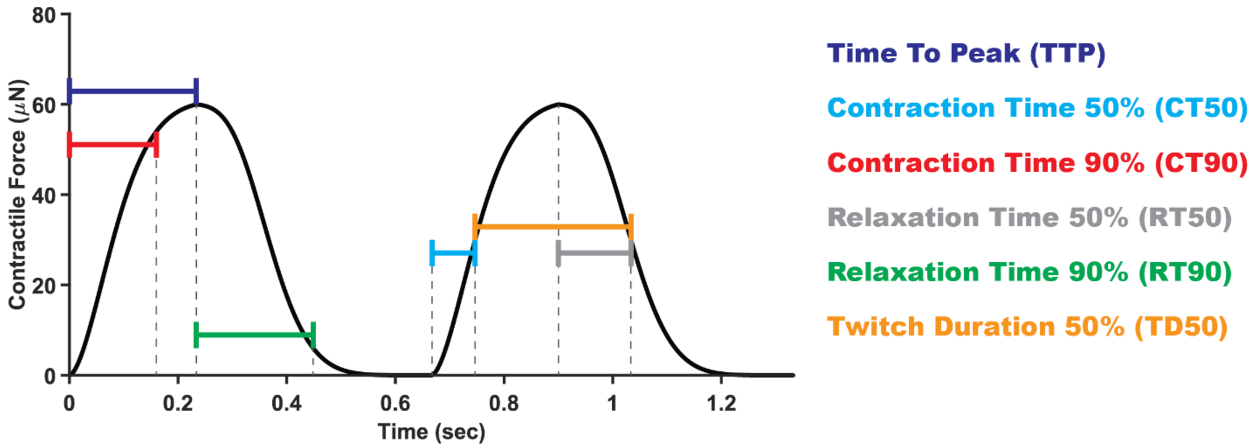


Figure S6. Schematic definition of twitch-timing metrics used to assess changes in twitch kinetics. Representative contractile-force traces illustrating the six timing metrics extracted from each twitch: time to peak (TTP), defined as the time from twitch onset to peak force; contraction time 50% and 90% (CT50 and CT90), defined as the time from contraction onset to 50% and 90% of peak force, respectively; relaxation time 50% and 90% (RT50 and RT90), defined as the time from peak force to 50% and 90% relaxation, respectively; and twitch duration 50% (TD50), defined as the interval between 50% of peak force on the rising and falling phases. Dashed vertical lines indicate the corresponding feature time points used for metric calculation.
